# Predictable and stable epimutations induced during clonal propagation with embryonic transcription factors

**DOI:** 10.1101/2022.03.15.484412

**Authors:** Anjar Tri Wibowo, Javier Antunez-Sanchez, Alexander Dawson, Jonathan Price, Cathal Meehan, Travis Wrightsman, Maximillian Collenberg, Ilja Bezrukov, Claude Becker, Moussa Benhamed, Detlef Weigel, Jose Gutierrez-Marcos

## Abstract

Although clonal propagation is frequently used in commercial plant breeding and plant biotechnology programs because it minimizes genetic variation, it is not uncommon to observe clonal plants with stable phenotypic changes, a phenomenon known as somaclonal variation. Several studies have shown that epigenetic modifications induced during regeneration are associated with this newly acquired phenotypic variation. However, the factors that determine the extent of somaclonal variation and the molecular changes associated with it remain poorly understood. To address this gap in our knowledge, we compared clonally propagated *Arabidopsis thaliana* plants derived from somatic embryogenesis using two different embryonic transcription factors-*RWP-RK DOMAIN-CONTAINING 4 (RKD4)* or *LEAFY COTYLEDON2 (LEC2)* and from two epigenetically distinct tissues. We found that both the epi(genetic) status of explant and the regeneration protocol employed play critical roles in shaping the molecular and phenotypic state of clonal plants. Phenotypic variation of regenerated plants can be largely explained by the inheritance of tissue-specific DNA methylation imprints, which are associated with specific transcriptional and metabolic changes in sexual progeny of clonal plants. Moreover, regenerants from roots were particularly affected by the inheritance of epigenetic imprints, which resulted in increased accumulation of salicylic acid in leaves and accelerated plant senescence. Collectively, our data reveal pathways for targeted manipulation of phenotypic variation in clonal plants.

## Introduction

Many multicellular organisms can reproduce both sexually and asexually, but clonal reproduction is especially common in plants, where it is prevalent in ferns, mosses and many angiosperms ^1,2^. Most plant species can switch to asexual reproductive programs in response to environmental stimuli by forming specialized structures such as rhizomes, stolons and bulbils ^2^. Such flexibility in reproductive strategies is thought to be possible due to the high degree of plasticity and totipotency of plant cells. This property has been traditionally exploited for the clonal propagation and genomic manipulation of many economically important species ^3^. Clonal propagation can be achieved in different ways, from simple methods such as grafting and cuttings to more advanced methods such as tissue culture-induced regeneration using phytohormones or embryonic/meristematic transcription factors. In *Arabidopsis thaliana* (hereafter, Arabidopsis), somatic embryogenesis (SE) and plant regeneration can be initiated through the ectopic overexpression of embryonic transcription factors such as *AGAMOUS-LIKE15 (AGL15)* ^4^, *BABY BOOM (BBM)* ^5^, *LEAFY COTYLEDON2 (LEC2)* ^6^, *WUSCHEL* (*WUS*) ^7-9^ and *RWP-RK DOMAIN-CONTAINING 4 (RKD4)* ^10^.

A key characteristic of asexual propagation is that the individuals produced are genetically identical to their parents, except for a few spontaneous *de novo* mutations ^1^. However, clonal plants often display heritable phenotypic variation ^11,12^, known as somaclonal variation, which is linked to genetic ^13,14^ and epigenetic changes ^12,15,16^. In a previous study we found that Arabidopsis plants regenerated through the ectopic activation of an embryonic transcription factor, using above- and belowground organs, resulted in clonal plants that inherited molecular signatures, DNA methylation and gene expression profiles, characteristic of the tissue of origin ^12^. Whether this molecular and phenotypic variation is primarily caused by the direct regeneration of clonal individuals from different organs, or driven by the organ-specific activity of the embryonic transcription factor employed, has been unclear. To address this gap in our knowledge, we used Arabidopsis to generate clonal progeny from roots and leaves through the ectopic activation of two different embryonic transcription factors. We found that the organ employed for cloning makes indeed an important contribution, with root regenerants inheriting tissue-specific DNA methylation imprints and transcriptional states, independently of the regeneration methodology used. Collectively, our study has revealed that the molecular changes and phenotypic variation stably inherited in clonal plants are a consequence of tissue-specific imprints propagated during cloning. In natural clonal plant populations, this phenotypic diversity could contribute to the adaptation to recurrent environmental challenges.

### Phenotypic variation present in clonal plants is underpinned by tissue of origin

In a previous study, we identified heritable molecular and phenotypic changes in plants propagated through the induction of the embryonic transcription factor RKD4 ^12^. To define how this somaclonal variation is created, we used Arabidopsis accession Ws-2 to generate clonal progeny from leaf and root tissues using transgenic lines carrying transgenes that allow for controlled expression of *LEC2* and *RKD4*, two transcription factor genes necessary for the initiation of different stages of embryo development ^6,10^. To determine the stability of the phenotypes arising during cloning, we propagated seeds from each regenerated individual by selfing over three consecutive generations (G1-G3) under similar growth conditions, thus limiting possible effects caused by tissue culture. When we grew G2 and G3 clonal progeny, we noticed that the morphology of plants regenerated from roots (RO) was noticeably different from non-regenerated plants, while leaf regenerants (LO) were indistinguishable from the control plants (Fig. 1A-B). The growth characteristic of RO-LEC2 plants resembled those found in regenerants created in Col-0 using RKD4 ^12^. Notably, most RO-LEC2 lines (4/5) had reduced biomass, smaller leaf area and senesced early (Fig. 1C-D), with different growth conditions having no discernible effects on these phenotypes (Fig. 1 Supplement 1). Collectively, our data suggest that the major factor contributing to phenotypic diversity in clonal plants is the tissue of origin employed for regeneration, thus pointing to the long-term inheritance of somatic epigenetic states in the sexual progeny of clonal plants.

**Fig. 1.**
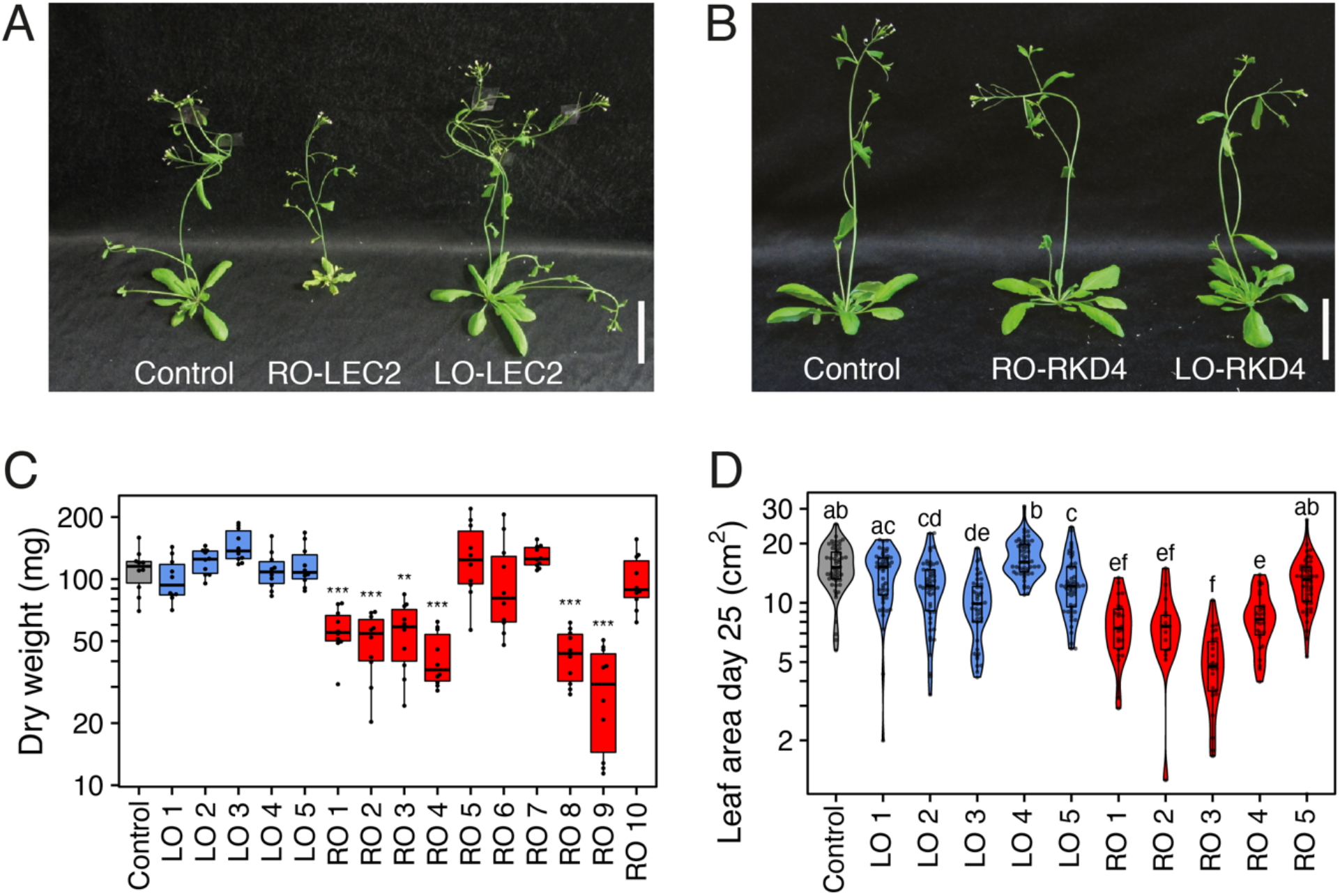
Growth phenotypes of Arabidopsis plants cloned from roots (RO) and leaves (LO) through LEC2/RKD4-induced regeneration and propagated by self-fertilization over three generations. (A) Representative images of 28 day old Arabidopsis Ws-2 plants. Control (Ws-2/indLEC2), and clonal RO-LEC2 and LO-LEC2 G3 progeny. Scale bars, 5 cm. (B) Representative images of 28-day old Arabidopsis Ws-2 plants. Control (Ws-2/indRKD4), and clonal RO-RKD4 and LO-RKD4 G3 progeny. Scale bars, 5 cm. (C) Boxplot showing variation in plant biomass for control (indLEC2/Ws-2) and independent G3 lines. Asterisks represent significant differences, determined by Student’s *t-test*, ** p<0.01, *** p<0.001, sample size n = 10. (D) Violin plot and boxplot of leaf area for control (Ws-2/indLEC2) and independent G3 lines. Letters represent groups of statistically significantly different samples (p<0.05), determined by Tukey’s test, sample size n = 20.

**Fig. 1 Supplement 1:**
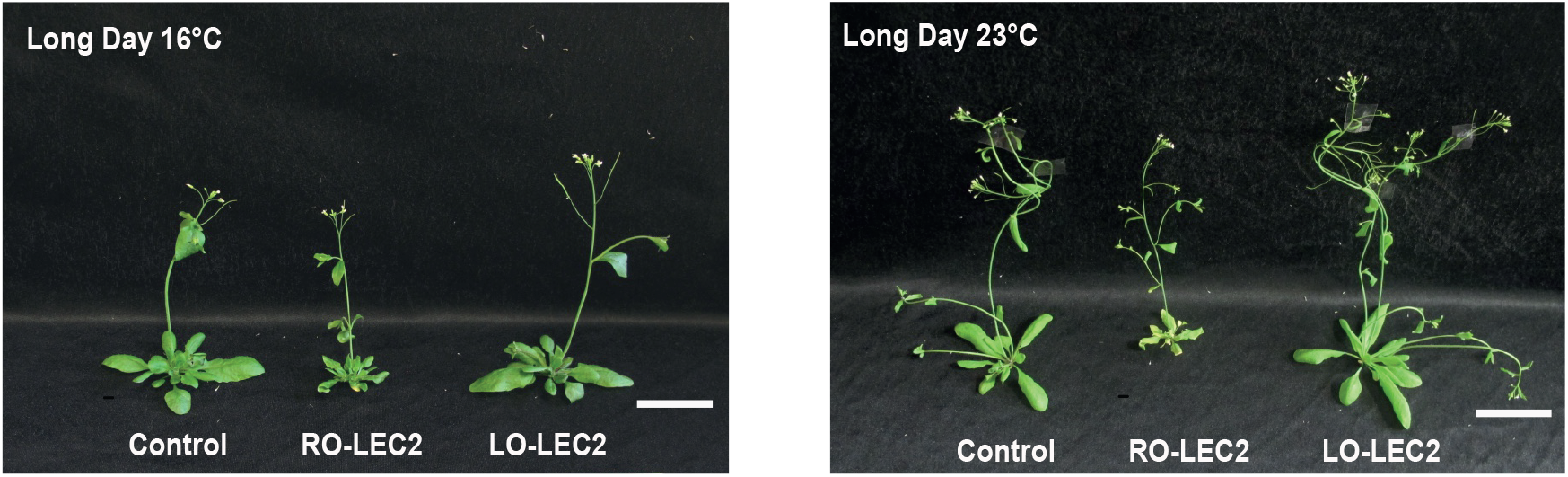
Arabidopsis Ws-2 plants from clonal RO-LEC2 and LO-LEC2 G3 progeny grown under different growth conditions.

### Tissue of origin affects the activity of defense-related genes

To determine what molecular changes might underpin the observed morphological variation, we performed whole-genome transcriptome analysis in leaves and roots from randomly selected, independent G3 lines. Using stringent thresholds [false discovery rate (FDR) <0.05; absolute log-two fold change >1.5], we obtained lists of Differentially Expressed Genes (DEGs) from comparing leaves and roots of regenerants and control plants (Supplementary tables S1.1 (RKD4) and S1.2 (LEC2)). There were notable differences between the regenerants induced by the two embryonic transcription factors; leaves of RKD4-induced regenerants were the most affected independently of the tissue of origin (LO or RO). Notably, we found more DEGs for LO than for RO plants (Fig. 2A and Fig. 2 Supplement 1), which resembled the effects found in Col-0 plants cloned using RKD4 ^12^. LEC2 regenerants had the highest numbers of DEGs in the opposite tissue from which the plant was regenerated, i.e., in roots of LO-LEC2 plants and in leaves of RO-LEC2 plants (Fig. 2B and Fig. 2 Supplement 2). Since gene expression was strongly dysregulated in the leaves of both RO-RKD4 and RO-LEC2 regenerants, we looked for an overlap between the DEGs. We found more shared dysregulated genes than expected by chance (p-value = 2.1×10^−17^), suggesting common effects independently of the regeneration strategy.

**Fig. 2.**
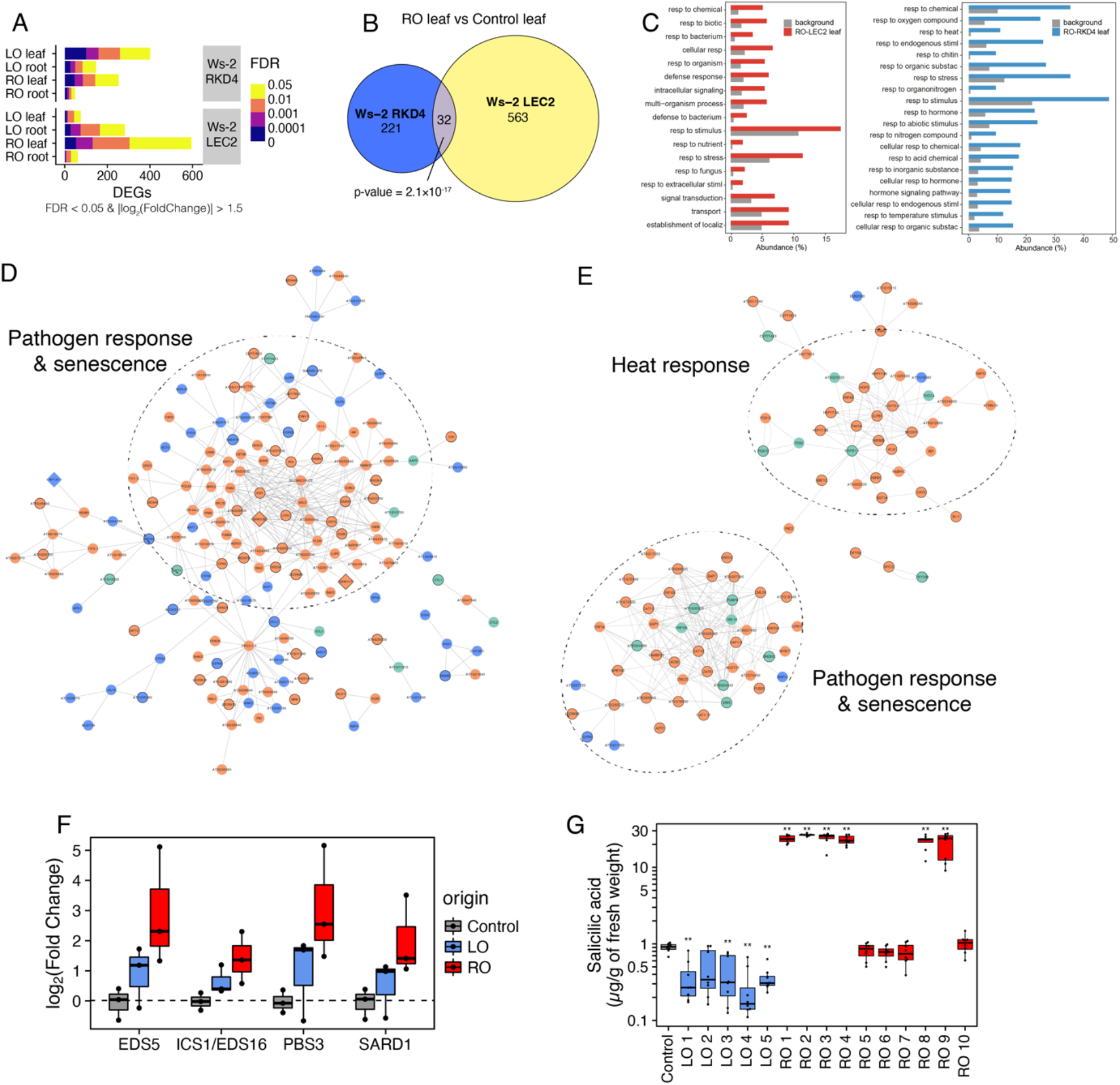
Differential expression of pathogen response and senescence genes in RO and LO plants. (A) Number of DEGs for different comparisons split by FDR levels. Data from three randomly selected regenerants for each tissue. (B) Euler diagram of common DEGs found in RO plants regenerated using two different transcription factors. p-value indicates results from Fisher’s exact test. (C) Gene Ontology analysis showing significant enrichment for stress-related genes among DEGs in RO-LEC2 (left) and RO-RKD4 (right) leaves. Categories are sorted by decreasing FDR. (D) Interaction network of DEGs found in leaves of RO-LEC2 plants. Orange, upregulated genes; blue, downregulated genes; green, non-DEGs with significant interactions with the network. Diamond shaped nodes, transcription factors. Main GO terms associated with clusters are highlighted. (E) Interaction network of DEGs found in leaves of RO-RKD4 plants. Orange, upregulated genes; blue, downregulated genes; green, non-DEGs with significant interactions with the network. Diamond shaped nodes, transcription factors. Main GO terms associated with clusters are highlighted. (F) Expression levels of genes associated with SA biosynthesis in LO-LEC2 and RO-LEC2 plants. WT, Ws-2/indLEC2 parent. (G) SA concentrations in leaves of LEC2 regenerants at the G3 generation. Asterisks represent significant differences, determined by Student’s *t-test*, ** p < 0.01, sample size of n = 10. Ws2 = Ws-2/indLEC2 parent.

To obtain further insight into the possible function of these DEGs, we performed Gene Ontology (GO) analyses, finding that DEGs in leaves from both RO-LEC2 and RO-RKD4 lines were enriched for stress and defense responses (FDR<0.05, Fig. 2C). In contrast, DEGs in roots from either LO-LEC2 or LO-RKD4 lines were primarily associated with metabolic processes (Fig. 2C). Network analysis revealed that genes related to pathogen responses and senescence were upregulated in RO plants, independently of the embryonic transcription factor used for clonal propagation (Fig. 2D-E). As several of the genes upregulated in RO-LEC2 leaves were part of the salicylic acid (SA) biosynthesis pathway (Fig. 2F), we measured the concentration of SA. Reduced biomass and early senescence in RO-LEC2 lines were strongly associated with high levels of SA, indicating a causal relationship between transcriptional changes, differences in production of a defense hormone known to negatively affect growth and to induce early senescence ^17^, and morphological changes in the regenerants (Fig. 2G). Collectively, our analyses revealed that the clonal propagation of plants using embryonic transcription factors results in the transmission of organ-specific transcriptional states.

**Fig. 2 Supplement 1:**
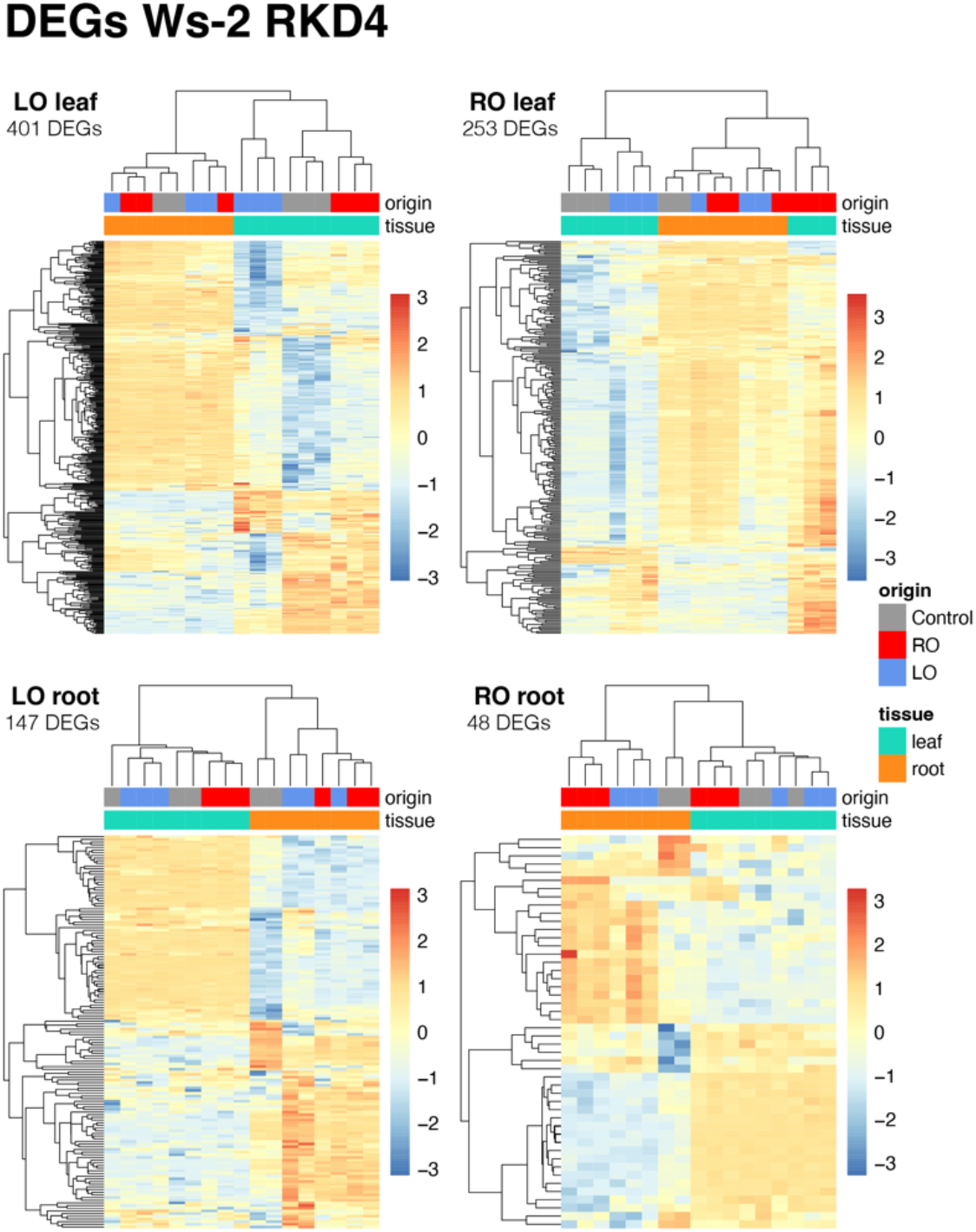
DEGs in LO-RKD4 and RO-RKD4 regenerants compared to Ws-2/indRKD4 control plants. Data from three randomly selected regenerants for each tissue.

**Fig. 2 Supplement 2:**
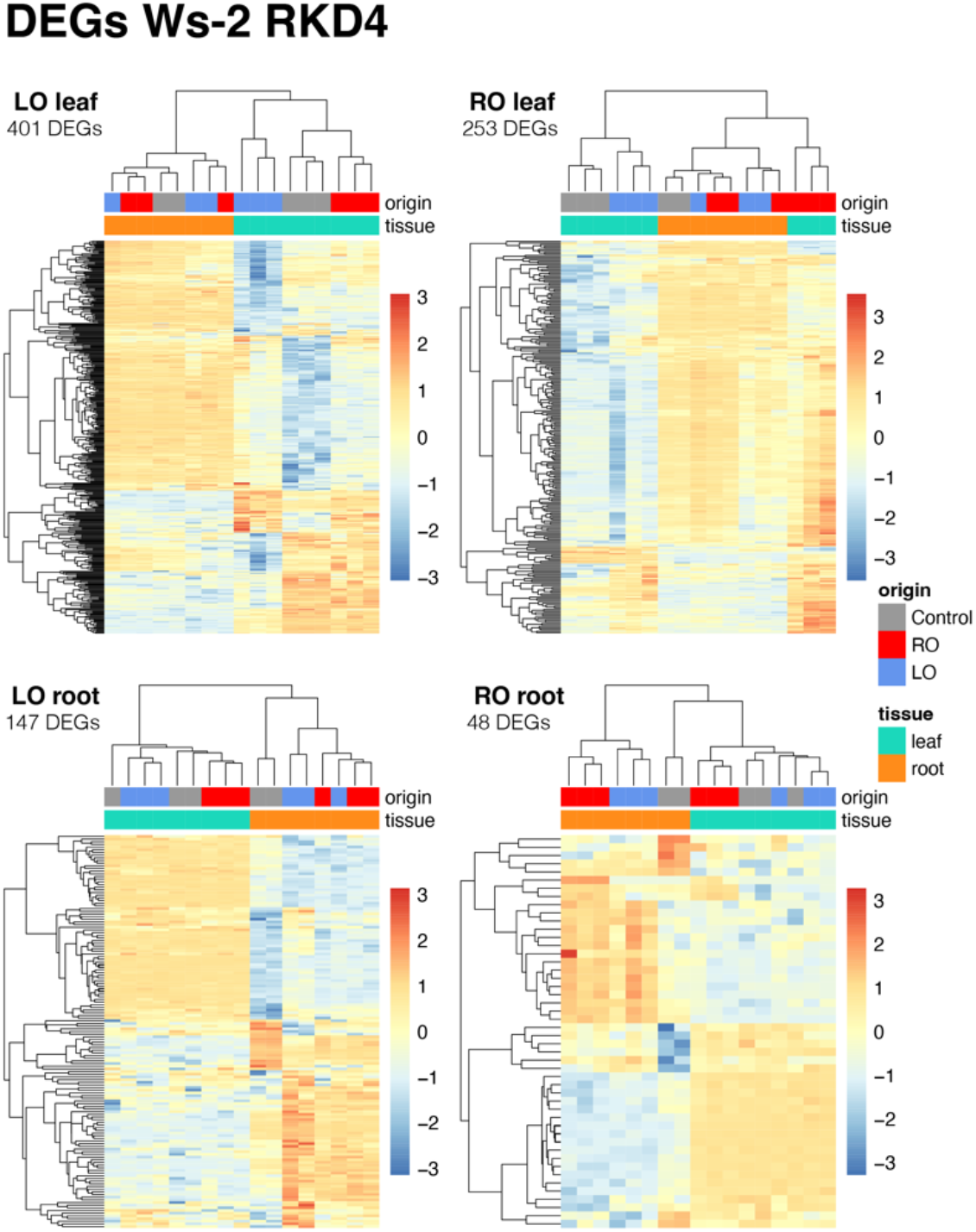
DEGs in LO-LEC2 and RO-LEC2 regenerants compared to Ws-2/indLECs control plants. Data from three randomly selected regenerants for each tissue.

#### Heritable DNA methylation imprints in LEC2-induced clonal progeny

We postulated that the phenotypic and transcriptional changes found in plants regenerated with embryonic transcription factors may be caused by the inheritance of heritable tissue-specific epigenetic imprints. To test this hypothesis, we studied the methylome of LEC2 regenerants that had been propagated for two successive generations (G2 and G3). To map methylome data we generated a *de novo* assembly of the Ws-2 genome (see methods). In an all-against-all comparison of RO-LEC2 and LO-LEC2 plants, we found 226, 120 and 386 Differentially Methylation Regions (DMRs) in CG, CHG and CHH contexts, respectively (Supplementary Table 2-List of DMRs). Principal component analysis (PCA) of the 226 CG-DMRs showed that the main source of variance in our dataset was the tissue analyzed (69% of variance), with methylome profile of leaves of RO-LEC2 plants being slightly skewed towards root samples (Fig. 3A). Another 6% of variance can be explained by the tissue of origin, separating RO-LEC2 and LO-LEC2 samples from each other and from the controls, independently of the tissue sampled. Some of the remaining variance (3%) comes from the regeneration process independently of tissue sampled or tissue used for cloning (Fig. 3A). These data suggest that even though the methylome of regenerated plants mainly corresponds to the characteristic methylome of the tissue sampled, a number of epigenetic imprints are maintained from the tissue employed for cloning, even after plants had undergone three cycles of sexual reproduction. Our data also show that regeneration by itself contributes to discrete methylation imprints that are stably inherited in clonal plants (Fig. 3 Supplement 1). Similar, albeit weaker patterns were observed for non-CG DMRs (Fig. 3 Supplement 1). To define the potential impact of these epigenetic changes, we mapped DMRs to genome features, which revealed that regions flanking protein coding genes were particularly likely to be associated with methylation changes (Fig. 3 Supplement 2).

**Fig. 3.**
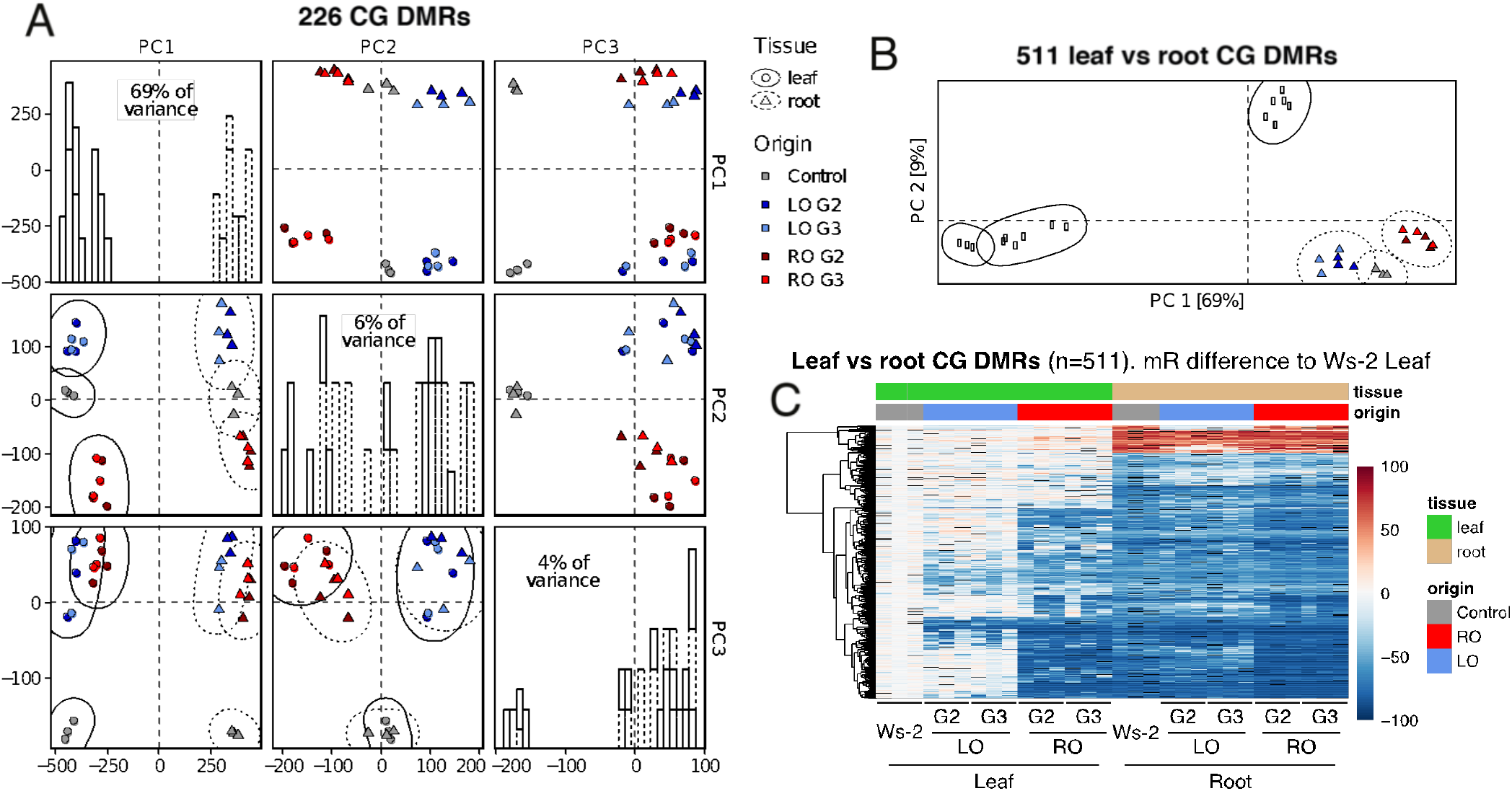
DNA methylomes of LEC2 regenerants. (A) PCAs of CG DNA methylation levels at 226 CG-DMRs identified in all-against-all comparison and histograms of the distribution of samples across each principal component. Probability ellipses (95% distribution) for each group are shown. Gray, control; Blue, LO-LEC2; Red, RO-LEC2. Data from three randomly selected regenerants for each tissue. (B) PCA of CG methylation levels at 511 CG-DMRs identified in the comparison of control leaves and roots. Probability ellipses (95% distribution) for each group are shown. Gray, control; Blue, LO-LEC2; Red, RO-LEC2. Data from three randomly selected regenerants for each tissue. (C) Heatmap of CG methylation levels in leaves and roots of control plants and LEC2 regenerants. Each column corresponds to an independently regenerated line. Ws-2 = Ws-2/indLEC2. Data from three randomly selected regenerants for each tissue.

When only considering DMRs distinguishing leaves and roots of non-regenerated plants, RO-LEC2 leaves, although distinct from other samples, resembled the methylation of root samples (Fig. 3B). Most DMRs in leaves of RO-LEC2 regenerants were primarily hypomethylated compared to leaves of control plants, resembling the methylation profile of these regions in control roots, and they were stably inherited (Fig. 3C and Fig. 3 Supplement 1). Similar to roots of control plants, these DMRs were also hypomethylated in roots of LO-LEC2 regenerants, thus pointing to active DNA demethylation in Arabidopsis roots. Collectively, our analyses demonstrate that induction of somatic embryogenesis and subsequent regeneration results in tissue-specific epigenetic imprints that are stable during sexual reproduction.

**Fig 3. Supplement 1.**
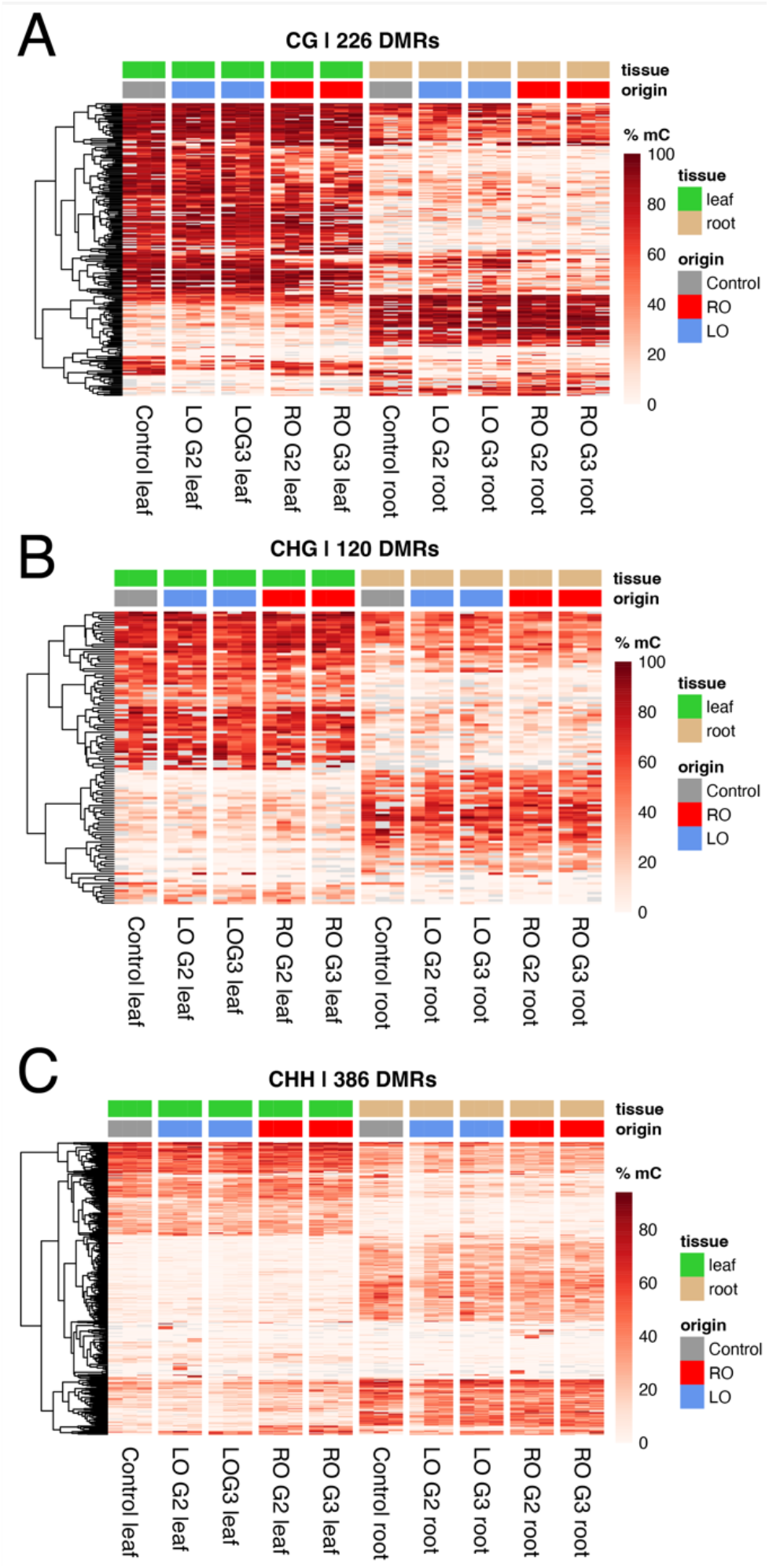
DNA methylation variation in LEC2 regenerants. (A) CG methylation levels at 226 CG-DMRs identified in an all-against-all comparison. (B) CHG methylation levels at 120 CHG-DMRs identified in an all-against-all comparison. (C) CHH methylation levels at 386 CHH-DMRs identified in an all-against-all comparison.

**Fig.3 Supplement 2.**
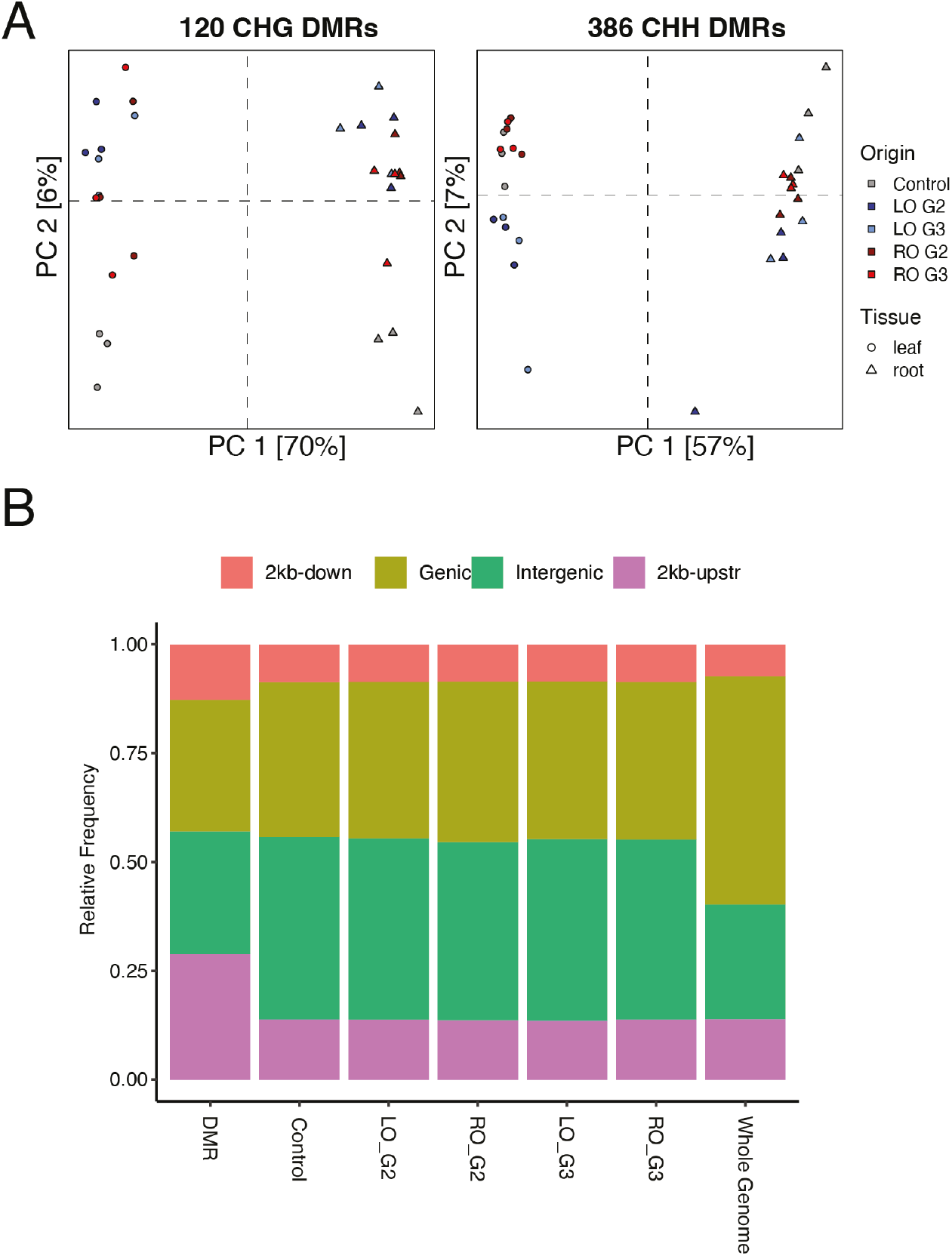
DNA methylation variation in LEC2 regenerants affects all contexts and appears mainly in proximity to genes. (A) PCAs of CHG DNA methylation levels at 120 CHG-DMRs identified in all-against-all comparison (left two panels) and PCAs of CHH methylation levels at 386 CHH-DMRs identified in all-against-all comparison (right two panels). Percentages in brackets indicate the proportion of variance explained by each principal component (PC). (B) Distribution of different genetic features in DMRs identified in all-against-all comparison for all three contexts, in methylated regions of different samples, and in the whole genome. Intergenic is defined as all sequences more than 2 kb away from a gene.

#### Precocious leaf senescence in root-regenerants caused by epigenetic deregulation of the salicylic acid pathway

To determine the molecular basis for the early senescence phenotype observed in RO regenerants, we performed reciprocal crosses between Ws-2 and RO-LEC2 plants. Five reciprocal F_1_ hybrids were self-pollinated to generate F_2_ seed progeny (Fig. 4A). In the F_2_ generation, some plants were early senescing, characteristic of RO-LEC2 plants, while others were not. The fraction of early-senescing plants was around one quarter, with 16% (13 in 80) early-senescing F_2_ progeny derived from Ws-2/indLEC2 x RO-LEC2 hybrids, and 23% (28 in 120) early-senescing F_2_ progeny derived from RO-LEC2 x Ws-2/indLEC2 hybrids (p>0.05, Fisher’s Exact test). These data suggest that early senescence in RO-LEC2 regenerants may be under the control of a single recessively acting locus. To determine whether this was due to a genetic mutation, we whole-genome shotgun sequenced 10 phenotypically normal and 10 early-senescing individuals from five different segregating F_2_ populations (Fig. 4A). We could not identify genetic variants linked to this phenotype (Fig. 4 Supplement 1), suggesting that the phenotypes were caused by an epigenetic change.

**Fig. 4.**
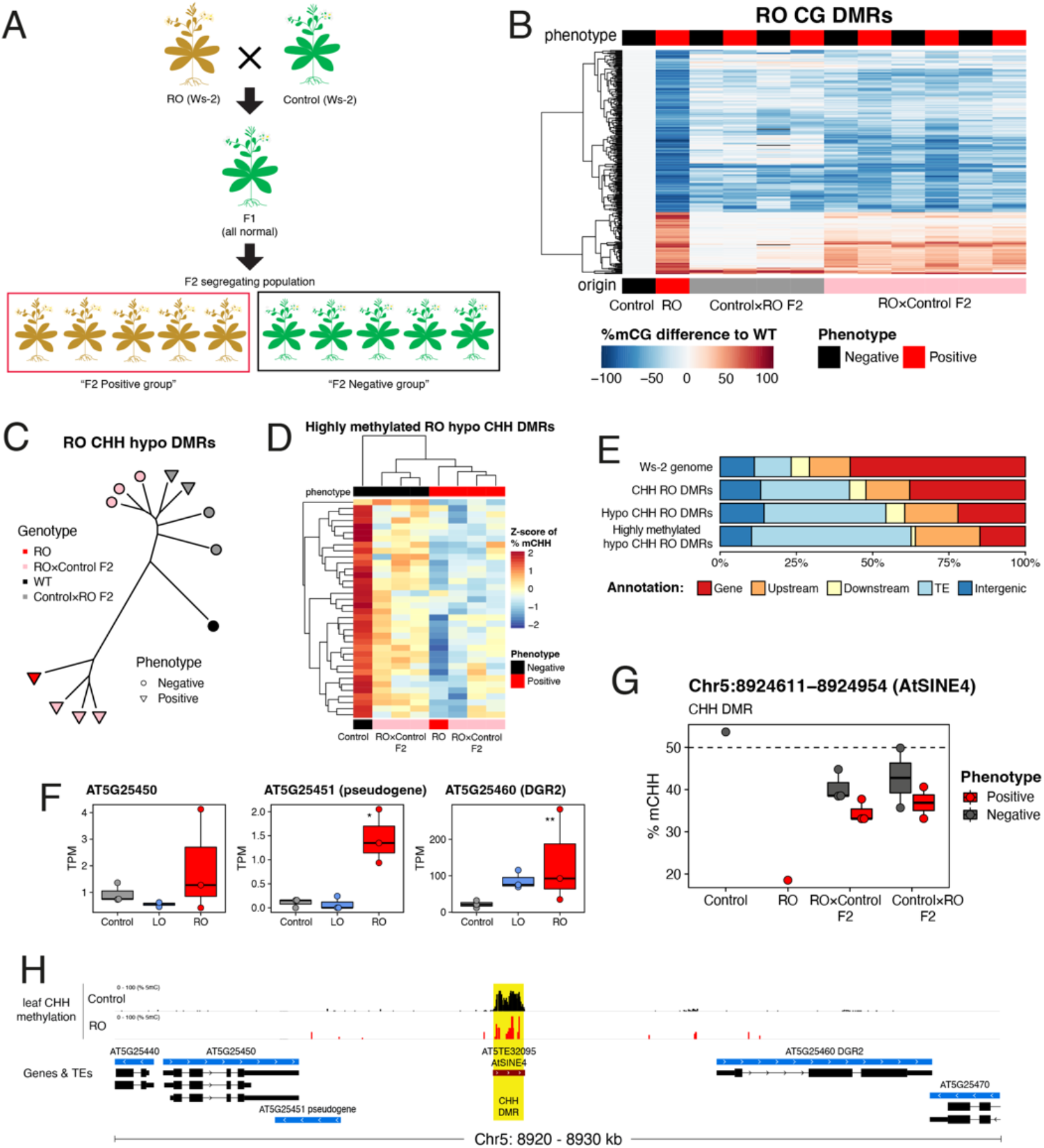
Leaf senescence and TE hypomethylation in RO-LEC2 root regenerants. (A) Experimental design to identify epi- and DNA mutations linked to RO-LEC2 phenotypes. Green indicates phenotypically normal plants (n=10) and brown early senescing ones (n=10). Analysis was carried out using five independent F_2_ progeny. (B) Heatmap showing gains and losses of CG methylation for RO-LEC2-leaf DMRs in F_2_ progeny from reciprocal backcrosses between control (Ws-2/indLEC2) and RO-LEC2 plants. Each column represents an independent F_2_ progeny individual. (C) Dendrogram of F_2_ progeny and parents at parental CHH-hypo-DMRs. (D) Heatmap of Z-scores in CHH methylation for highly methylated (Ws-2/indLEC2 %mC > 50) parental CHH-DMRs, showing clustering of parents and RO x Ws-2/indLEC2 F_2_ progeny. (E) Distribution across features of different subsets of parental CHH DMRs. (F) Expression levels for genes proximal to Chr5:892411-894943 (AtSINE4) DMR in leaves of control (Ws-2/indLEC2) and regenerants. (G) CHH methylation levels for Chr5:892411-894943 (AtSINE4) DMR in Ws-2/indLEC2, RO-LEC2, and F_2_ reciprocal crosses, showing association between demethylation and morphological phenotype. (H) Genomic view around Chr5:892411-894943 (AtSINE4) DMR. Yellow box represents RO-LEC2 CHH-DMR.

We investigated the mode of inheritance of RO imprints by generating DNA methylation profiles of leaves from F_2_ progeny derived from Ws-2/indLEC2 x RO-LEC2 hybrids. If differences in DNA methylation were the cause for the growth phenotypes characteristic of RO plants, the methylation imprints must be already present in the parents. We therefore focused on DMRs between leaves of Ws-2/indLEC2 and RO-LEC2 plants. Most regions that are hypomethylated in RO-LEC2 regenerants were stably inherited both mitotically and meiotically (Fig. 4B). On the other hand, most hypermethylated DMRs had disappeared after two generations of paternal transmission, although they were faithfully inherited maternally (Fig. 4A). This was observed for both CG- and CHG-hypermethylated DMRs, while CHH-hypermethylation was reset both paternally and maternally (Fig. 4 Supplement 2). These data suggest that the early senescence phenotype observed in RO-LEC2 plants may be caused by an epimutation associated with a reduction in DNA methylation, as plants with early senescence appeared in both directions of the cross. After clustering samples based on the levels of methylation for DMRs, we found that CHH-hypomethylated DMRs as a class predicted phenotype, (Fig. 4C), and this was clearest for RO-LEC2 x Ws-2/indLEC2 progeny. Thirty-six DMRs co-segregated with the growth phenotype. Normal plants were highly methylated, as in the Ws-2/indLEC2 control, and early senescing plants had low methylation at these 36 DMRs (Fig. 4D). These phenotype-associated DMRs were strongly enriched in transposable elements (TEs) (Fig. 4E). Collectively, these data suggest that demethylation of TEs is associated with early senescence of RO-LEC2 plants.

To understand how losses of DNA methylation at specific loci could cause the phenotype, we searched for deregulated genes in the proximity of the 36 DMRs. We found two genomic regions where methylation status co-segregated with the phenotype for both directions of the crosses (Fig. 4G [will change to F]). One region included three genes in its proximity that are upregulated in leaves of RO-LEC2 regenerants (Fig. 4F [will change to G]). The DMR is located in a TE of the AtSINE4 family, between two of the genes found to be misregulated in RO-LEC2 plants when compared to Ws-2/indLEC2 control (Fig. 4H). We investigated the chromatin conformation of these regions in leaves and roots of wild-type plants, and found notable differences, thus suggesting it may act as a long-distance regulatory element for flanking genes (Fig. 4 Supplement 3). Another DMR associated with early senescence in RO-LEC2 plants was located in an AtMUNX1 TE upstream of the *PRE1/BNQ1* locus (Fig. 4 Supplement 4), in a region flanking a CarG box known to be a target of MADS box containing transcription factors ^18^ and negatively regulated by the transcription factor pair APETALA3 (AP3) and PISTILLATA (PI) ^19^.

**Fig. 4 Supplement 1.**
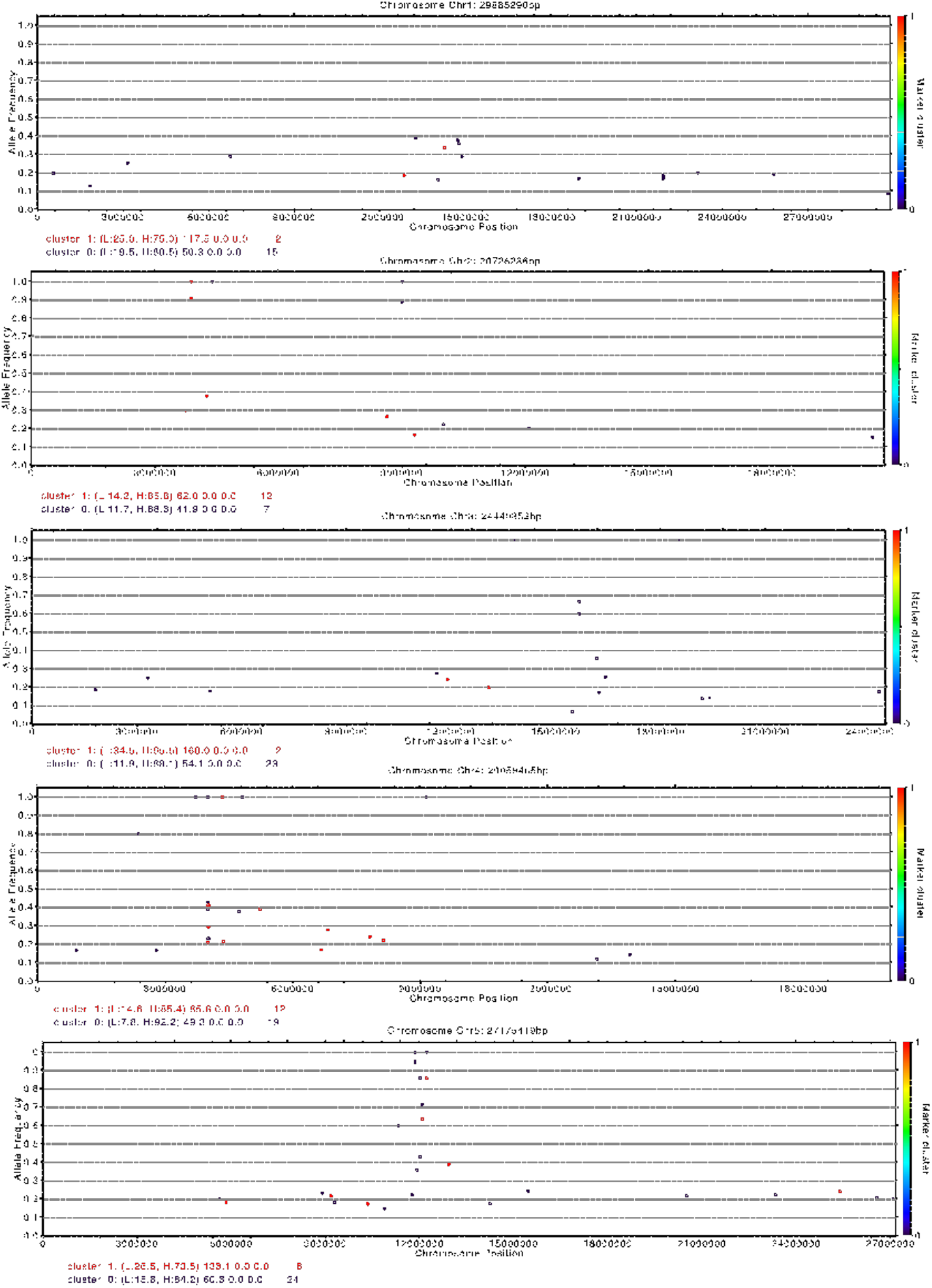
Distribution of SNPs associated with early senescence of RO-LEC2 plants. Allelic frequencies across chromosomes of foreground markers derived from comparing phenotypically positive (foreground) against phenotypically negative (background) plants.

**Fig. 4 Supplement 2.**
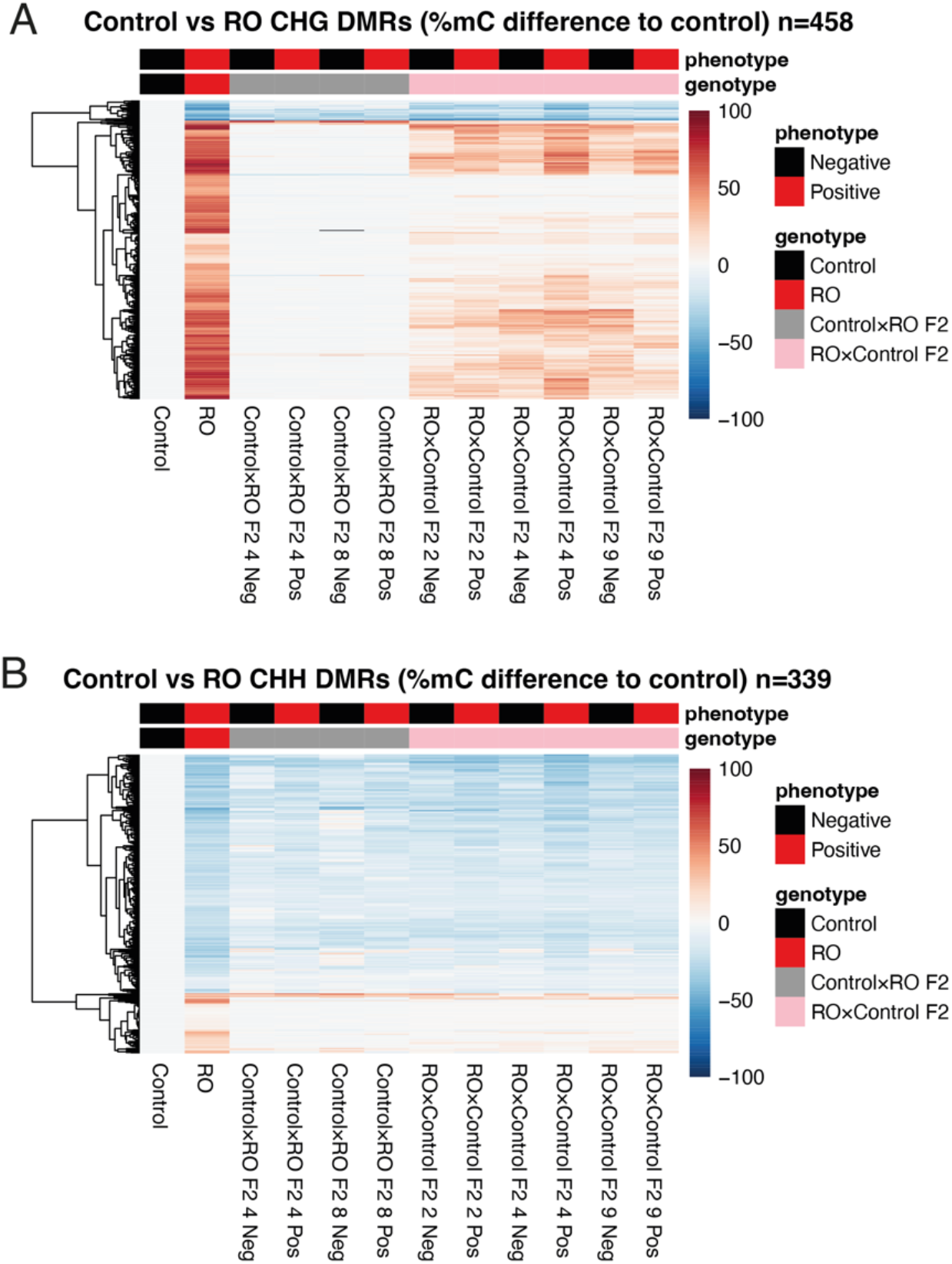
Hypermethylated RO-LEC2 DMRs revert to normal methylation levels after backcrossing. Heatmaps showing gains and losses of CHG (A) and CHH (B) DNA methylation levels for RO-LEC2 leaf DMRs in F_2_ progeny from reciprocal backcrosses between control (Ws-2/indLEC2) and RO-LEC2 plants. Each column represents plants from independent F_2_ progenies.

**Fig. 4 Supplement 3.**
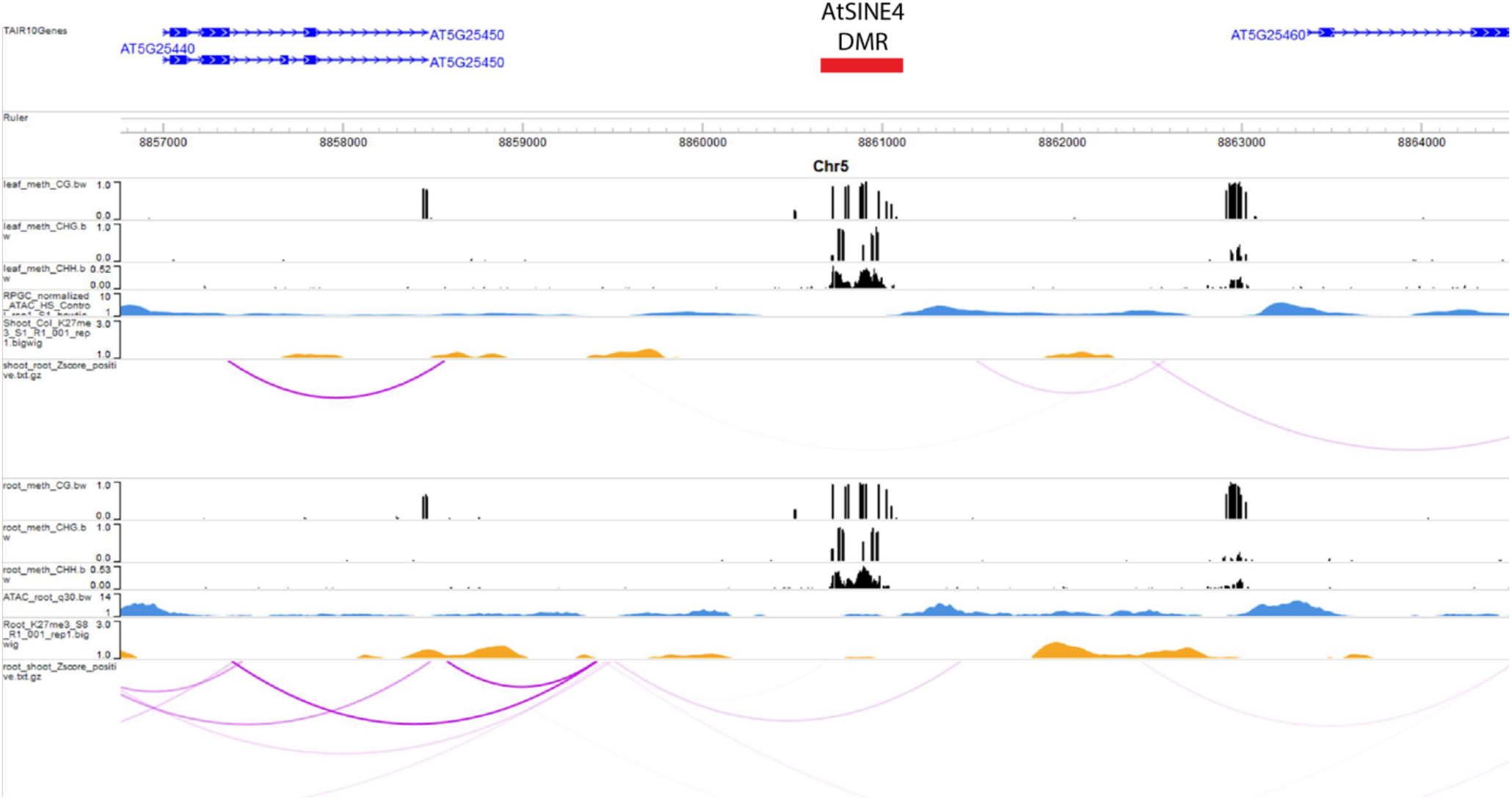
Epigenome browser view of AtSIN4 genomic region showing DNA methylation, ATAC-seq, H3K27me3 and Hi-C data from roots and shoots.

**Fig. 4 Supplement 4.**
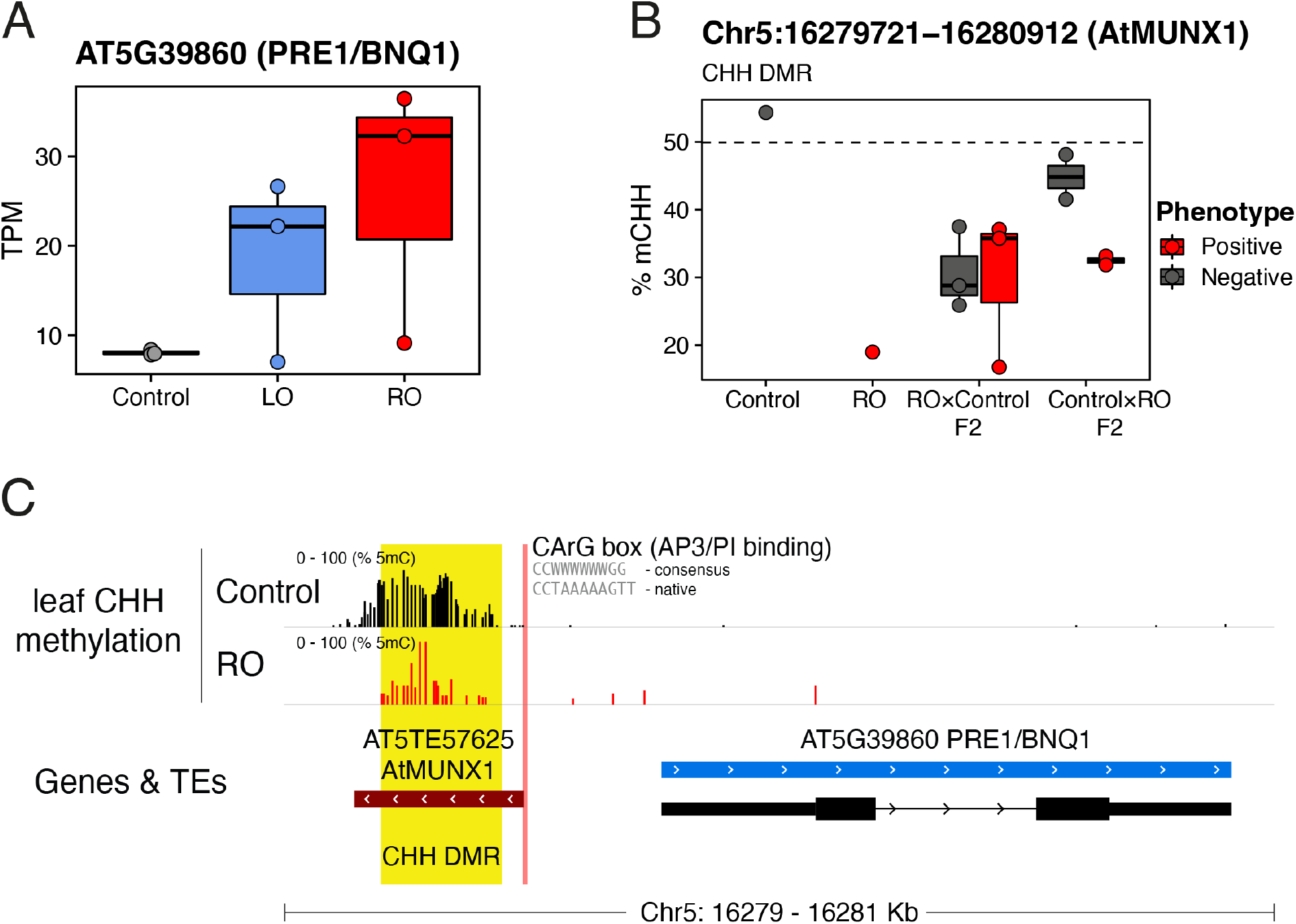
Expression of *PRE1/BNQ1* is associated with the hypomethylation of a flanking AtMUNX1 transposon.

## DISCUSSION

Previous studies have shown that clonal plants regenerated through tissue culture can have new phenotypes that may be an indirect consequence of the methodology employed for regeneration. For instance, plant regeneration using phytohormones induces stochastic changes in DNA methylation, primarily hypomethylation, that are associated with changes in gene expression ^11,13,16,20,21^. On the other hand, regeneration after the induction of somatic embryos using embryonic transcription factors results in nonrandom changes in DNA methylation, which are heritable and associated with transcriptional and phenotypic variation ^12^. To define the factor(s) that may contribute to the molecular and phenotypic diversity found in plants propagated through somatic embryogenesis, we conducted a detailed analysis on regenerants created by the ectopic expression of LEC2 and RKD4, two transcription factors that are part of distinct embryonic transcriptional networks ^10,20,22^. We found tissue-specific epigenetic imprints in these regenerants that were associated with gene expression and phenotypic variation.

The molecular mechanism(s) underpinning the inheritance of the tissue-specific epigenetic imprints that accumulate in these clonal plants are yet unknown, but they may be linked to the capacity of these two transcription factors to reprogram the development of somatic cells to adopt an embryonic program without altering their epigenomic configuration. We postulate that the tissue-specific imprints arising during cloning may be developmentally programmed, since leaves and roots have distinct epigenetic characteristics ^23-27^. The tissue-specific imprints found in clonal plants may be linked to the distinct chromatin architecture found in root and shoot cells, which is largely influenced by chromatin loops associated with histone 3 lysine 27 trimethylation ^28^. Chromatin interactions likely contribute to the developmental differentiation of specific cell types in plants, as it is the case in animals ^29,30^, and may contribute to the stability of epigenetic imprints in clonal plants.

In our system, plants regenerated from roots senesced early, a phenotype that was inherited by sexual progeny either after selfing or outcrossing. This early leaf senescence strongly correlated with DNA hypomethylation of specific TEs flanking genes associated with the SA pathway. Several studies have shown that the long-distance regulatory action of TEs in Arabidopsis is associated with changes in DNA methylation ^31^. Moreover, transposon mobilization and variation in TE methylation have been linked to environmental adaptation in Arabidopsis populations ^32^. Our data suggest that epimutations at TEs could accumulate in populations of species that reproduce asexually, and that these epimutations could confer an adaptive advantage under fluctuating environments.

The SA pathway may be an important target for the accumulation of epimutations as it si critical for plant defense ^33^ and it is strongly correlated with plant biomass, a characteristic that has traditionally been selected by breeders to increase yield in various crops, including maize ^34^, rice ^35^, and wheat ^36^. Notably, mutants that cause a global change of chromatin organization, such as *CROWDED NUCLEI* (*CRWN*), display constitutive activity of the salicylic acid-dependent pathway of immunity and *crwn* plants display dwarfism and premature senescence ^37,38^.

Epigenetic imprints generated during cloning can be inherited over multiple cycles of sexual reproduction, suggesting that the epigenetic machinery in plants is unable to repair these epimutations ^11,12,16,22^. That stable epimutations accumulate has been also documented in crosses between wild-type plants and mutants defective in DNA methylation ^39-43^ and histone demethylation ^44^. Notably, the tissue-specific DNA hypomethylation imprints arising in plants cloned using ectopic induction of LEC2 are transmitted through both the maternal and paternal germline to the offspring. However, DNA hypermethylation imprints in these plants are transmitted only maternally. The molecular mechanisms implicated in the resetting of these DNA hypermethylated imprints are not yet known but they may be linked to an active epigenomic reprogramming taking place during male gametogenesis ^45,46^ or embryo development ^47^.

In summary, our study has revealed that clonal propagation using embryonic transcription factors enables the generation of specific epigenomic changes to engineer novel phenotypes. Our data indicate that these epigenomic changes are stable and could be integrated in breeding programs to enhance the potential of wild and domesticated crop species.

## Acknowledgments

We thank Gary Grant for help with plant husbandry, and Liliana M. Costa for discussions and comments on the manuscript. Supported by ERA-CAPS AUREATE (DFG), and the Max Planck Society (D.W.), a European Research Council (ERC) Marie Skłodowska-Curie Fellowship (751204-H2020-MSCA-IF-2016) and and Airlangga University Hibah Riset Mandat Grant (387/UN3.14/PT/2020) (A.T.W.), and BBSRC grants (BB/L003023/1, BB/N005279/1, BB/N00194X/1 and BB/P02601X/1) (J.G-M.).

## Author contribution

A.T.W., D.W. and J.G.-M. conceived the project. A.T.W., J.A.S, and C.B conducted experiments. A.T.W., J.A.S, A.D, J.P., C.M, T.W., M.C., M.B. and I.B. and analyzed the data. D.W. and J.G.-M. supervised the research. A.T.W., J.A.S, D.W. and J.G.-M. wrote the manuscript with help from all other authors.

## Data and Materials Availability

Sequence data (BS-seq, RNA-seq and DNA-seq) have been deposited at the European Nucleotide Archive (ENA).

## METHODS

### Plant growth and clonal propagation

For all experiments, *A. thaliana* Ws-2 plants were grown in long day conditions (16-h light/8-h dark, 22 °C). To facilitate direct plant regeneration from different type of organs we used transgenic lines harboring an chemically-inducible transgenes to overexpress the GRANDE (GRD)/RWPRK motif-containing (RKD4) transcription factor ^10^ (indRKD4) or the B3 domain transcription factor LEAFY COTYLEDON2 (LEC2) ^48^ (indLEC2). Seeds from transgenic indRKD4 and indLEC2 lines were germinated on Murashige and Skoog (MS) medium containing 30 μM dexamethasone or 50 μM estradiol, respectively, and incubated for 14 days. Plants were transferred to MS media without dexamethasone or estradiol for 7 days to allow the formation of somatic embryos, which were then isolated and transferred to fresh MS plates using tungsten needles. After 2-3 weeks, the somatic embryos from leaves (LO) or roots (RO) were fully regenerated into new plants. We generated 10 independent regenerants from each organ (G0 generation), which were grown in soil to produce seeds. We grew 24 plants from each line (G1 generation) and selected 10 individuals at random for each regenerated line to produce seed. We continued the sexual propagation of each regenerant population for two additional generations (G2 and G3) following the same scheme.

### Plasmid constructs and transgenic lines

To generate estradiol-inducible RKD4 lines (indRKD4) we cloned using Gateway recombination the coding region of GRD/RKD4 (AT5G53040) in pMDC7 ^49^ modified by the insertion after PmeI digestion of a pOLE1:OLE1-RFP fragment from pFAST-R ^50^. The estradiol-inducible LEC2 lines (indLEC2) were generated by amplifying the LEC2 coding sequence by PCR and cloning in pBI-ΔGR vector. The LEC2-GR fragment was then cloned in pCGN18 ^51^ to create p35S::LEC2-GR. Plants were transformed by floral dipping, transgenic plants were selected in Kanamycin (indLEC2) or RFP seed selection (indRKD4), and propagated for three generations to isolate homozygous lines.

### Phenotypic analysis

For biomass analysis, plants were grown under long days for 4 weeks (16-h light/8-h dark, 22°C). Dry weight was recorded using aerial parts excised and dried at 80°C for 48 hours. For image-based phenotyping, images were acquired once per day in top view using two cameras per tray. The cameras were equipped with OmniVision OV5647 sensors with a resolution of 5 megapixels. Each camera was controlled by a Raspberry Pi computer (Revision 1.2, Raspberry Pi Foundation, UK). Stitched whole-tray images were separated into individual pots using a predefined mask for each position. Segmentation of plant tissue against background was performed by removing the background pixels and applying GrabCut-based automatic post processing. The leaf area of each plant was calculated based on the segmented plant images.

### SA measurement

The total levels of SA and SA glucoside (SAG) were measured from each sample using the *Acinetobacter* sp. ADPWH-*lux* biosensor ^52^. In brief, 175 mg of leaves material were harvested from four-week-old plants and homogenized in 250 μl 0.1 M sodium acetate buffer pH 5.5. Samples were then centrifuged for 15 min at 16,000 g. For total SA measurement, 200 μl of supernatant was transferred to a new tube and incubated with 20 μl of 0.5 U/μl β-glucosidase (Sigma-Aldrich) for 90 min at 37°C. Following β-glucosidase treatment, 60 μl of LB, 30 μl of plant extract, and 50 μl of *Acinetobacter* sp. ADPWH-*lux* (OD = 0.4) were incubated together at 37°C for 150 min. Luminescence was read with a Tecan luminometer. SA levels were calculated based on the SA standard curve, which was constructed using known amounts of SA stock (from 0 to 2500 ng/μl) diluted in *NahG* plant extract ^53^.

### Genome assembly and annotation

Plants were grown in long day conditions (16 h of light) at 23°C and a relative humidity of 65%. Philips GreenPower TLED modules (Philips Lighting GmbH, Hamburg, Germany) with 110-140 μmol m^-2^ s^-1^ light were used. Plants were grown for 3 weeks. In order to reduce the accumulation of starch all plants were placed in darkness for 30 h before harvesting young leaves. Fresh plant material was immediately flash-frozen after harvesting. HMW DNA extraction was performed following a custom protocol. Approximately thirty grams of leaf tissue were ground in liquid nitrogen and subsequently transferred into a 1 L flask containing 500 mL of ice-cold nuclei isolation buffer. The NIB was composed of 10 mM Tris (pH 8), 0.1 M KCl, 10 mM EDTA (pH 8), 0.5 M sucrose, 4 mM spermidine, and 1 mM spermine-4HCl. Flasks with ground plant material and NIB were kept on ice for fifteen minutes with occasional gentle swirling. The homogenate was filtered through four layers of miracloth (Merck, Germany). Subsequently, NIB solution with 20 % Triton X-100 was added. The solution was kept on ice for fifteen minutes with occasional gentle swirling. Afterwards, the solution was distributed into 50 mL Falcon tubes and centrifuged for 15 min at 3,250 rpm at four degrees using a tabletop centrifuge. Nuclei containing pellets were washed by adding a 40 mL NIB solution with 1 % Triton X-100 prior to centrifugation for 15 min at 4°C and 3,250 rpm. Subsequently, the supernatant was discarded before resuspending the nuclei pellet in 30 mL prewarmed (37 C) G2 lysis buffer (Qiagen, Germany). Resuspended pellets were incubated with RNase A (50 μg/ml) for 30 minutes while gently inverting the tube every five minutes. Proteinase K was added (250 μg/ml) and samples were incubated overnight at 50°C. Subsequently, the lysate was centrifuged at 8,000 rpm at 4°C for 15 min. The supernatant was poured onto an equilibrated Genomic Tip 100 (Qiagen, Germany). Afterwards the Genomic Tip 100 was washed twice with 7.5 mL QC Buffer (Qiagen, Germany) per wash. Genomic DNA was eluted from the column using 5 mL of prewarmed (50°C) QF Buffer (Qiagen, Germany). DNA was precipitated by adding 0.7 volumes of isopropanol. HMW DNA was obtained by spooling DNA strings onto a glass hook. The spooled DNA pellet was washed with 80% EtOH prior to resuspending it in 300 μL EB. DNA concentration was measured using Qubit HS DNA kit.

For library construction, high molecular weight DNA was sheared with a Megaruptor (Diagenode) set to a 40 kb fragment size. The DNA library was prepared using the Accel-NGS XL Library Kit (Swift Biosciences, United States). The library was size-selected with a minimum fragment size of 20 kb using a BluePippin (Sage Science, Beverly, United States and a High-Pass v3 cassette. The library was sequenced on the Pacific Biosciences (Menlo Park, United States) Sequel platform (ICS v5.0.0) with the v5.0.1 chemistry, which produced 5.09 Gb of long read sequences with a subread N50 of 15,750 bp.

The genome was assembled into 120 contigs of total length of 123.5 Mb with an N50 of 6.57 Mb using a custom Snakemake pipeline, “auto-asm” (https://github.com/weigelworld/auto-asm). Briefly, long reads were assembled into contigs using canu ^54^ and scaffolded without polishing steps into chromosomes against TAIR10 using reveal ^55^. Whole Genome Annotation was lifted over using the CAT ^56^ pipeline with TAIR10 as the reference genome and annotation.

### Transcriptome analysis

For mRNA-seq analysis, total RNA (10 μg) for each sample was used to purify polyA+ mRNA that was used for synthesis and amplification of cDNA. The RNA-seq libraries were prepared using TruSeq RNA Sample Preparation Kit from Illumina (San Diego, CA). RNA libraries were sequenced on an Illumina HiSeq2000 instrument (100 bp single-end). Read quality was assessed using FastQC and trimming of low-quality bases at the 3’ end of reads and adapter removal was done using Trimmomatic. Reads were mapped to the TAIR10 reference genome using Tophat (parameters -i 20 -I 30,000) with on average 85.4% unique mappings (min = 82.2, max = 88.8). The read counts for these libraries are given in Dataset S1. We used the R package DESEQ2 (version 1.10.1) ^8^. Genes were classified as significantly differentially expressed at either an FDR<0.01 or <0.05 and |log2 fold-change| >1.5. We used AgriGO v2.0 ^57^. The background used for identifying enrichment was the suggested background provided by AgriGo, which contains all GO annotated genes in Arabidopsis. Enrichment was performed using a hypergeometric test, p-values were corrected by multiple correction (Benjamini Hochberg FDR; α = 0.05).

### Identification of polymorphisms in RO-LEC2 lines

For library construction, high molecular weight DNA from individual plants was isolated using DNeasy Plant mini kit (Qiagen). The DNA library was prepared using a NxSeq AmpFREE Low DNA Library kit (Lucigen) and sequenced on a Illumina HiSeq2000 with a 100bp paired-end mode. Sequencing reads were pre-processed using fastp with default settings and the overrepresentation analysis parameter to remove reads with low quality score, irregular GC content, short length, and sequencing adapters present ^58^. Trimmed reads were then quality checked with outputs from fastp to ensure trimming had been successful. Trimmed reads were mapped to the TAIR10 reference genome using bowtie2 ^59^. Variant calling was performed on aligned and sorted BAM files using samtools mpileup and piped to bcftools to produce VCF files ^60^. VCF files were converted to SHOREmap format and analysed using SHOREmap backcross and then annotated to output tables and plots of backcross SNPs ^61^.

### Bisulfite sequencing

Bisulfite libraries were prepared from 100 ng genomic DNA using the TruSeq Nano kit (Illumina, San Diego, CA, USA) according to the manufacturer’s instructions, with the following modification. After adapter ligation and cleanup, samples were bisulfite-treated using the Epitect Plus kit (Qiagen, Hilden, Germany). Thermal cycle incubation was repeated once before clean-up. DNA fragments were amplified using the Kapa HiFi Hotstart Uracil+ PCR mix in 14 cycles of PCR (Kapa Biosystems, Wilmington, MA, USA). Bisulfite libraries were sequenced on an Illumina HiSeq3000 instrument (150 bp paired-end reads).

### Computational analysis of paired end BS-seq

Paired-end quality was assessed using FASTQC ^62^. Trimmomatic ^63^ was used for quality trimming. Parameters for read quality filtering were set as follows: Minimum length of 40 bp; sliding window trimming of 4 bp with required Phred quality score of 20. Trimmed reads were mapped to the *Arabidopsis thaliana* TAIR10 genome assembly using bwa-meth ^64^ with default parameters. Mapped reads were deduplicated using picardtools ^65^, and numbers of methylated/unmethylated reads per position were retrieved using MethylExtract ^66^. We calculate differentially methylated regions (DMRs) using MethylScore ^67^.

### Data visualization

For visualizing BS-seq genomic data we used Integrative Genomic Viewer (IGV) ^68^. For figures, we used R version 3.5.1 (www.r-project.org) with packages ggplot2 ^69^, eulerr ^70^ and pheatmap ^71^.

## REFERENCES

1 de Meeus, T., Prugnolle, F. & Agnew, P. Asexual reproduction: genetics and evolutionary aspects. Cell Mol Life Sci 64, 1355–1372, doi:10.1007/s00018-007-6515-2 (2007).

2 Zhang, Y. & Zhang, D. Asexual and sexual reproductive strategies in clonal plants. Frontiers of Biology in China 2, 256–262, doi:10.1007/s11515-007-0036-0 (2007).

3 Ikeuchi, M., Ogawa, Y., Iwase, A. & Sugimoto, K. Plant regeneration: cellular origins and molecular mechanisms. Development 143, 1442–1451, doi:10.1242/dev.134668 (2016).

4 Zheng, Q., Zheng, Y. & Perry, S. E. AGAMOUS-Like15 promotes somatic embryogenesis in Arabidopsis and soybean in part by the control of ethylene biosynthesis and response. Plant Physiol 161, 2113–2127, doi:10.1104/pp.113.216275 (2013).

5 Boutilier, K. et al. Ectopic expression of BABY BOOM triggers a conversion from vegetative to embryonic growth. Plant Cell 14, 1737–1749, doi:10.1105/tpc.001941 (2002).

6 Stone, S. L. et al. Arabidopsis LEAFY COTYLEDON2 induces maturation traits and auxin activity: Implications for somatic embryogenesis. Proc Natl Acad Sci U S A 105, 3151–3156, doi:10.1073/pnas.0712364105 (2008).

7 Bouchabke-Coussa, O. et al. Wuschel overexpression promotes somatic embryogenesis and induces organogenesis in cotton (Gossypium hirsutum L.) tissues cultured in vitro. Plant Cell Rep 32, 675–686, doi:10.1007/s00299-013-1402-9 (2013).

8 Love, M. I., Huber, W. & Anders, S. Moderated estimation of fold change and dispersion for RNA-seq data with DESeq2. Genome Biol 15, 550, doi:10.1186/s13059-014-0550-8 (2014).

9 Wang, K. et al. The gene TaWOX5 overcomes genotype dependency in wheat genetic transformation. Nat Plants 8, 110–117, doi:10.1038/s41477-021-01085-8 (2022).

10 Waki, T., Hiki, T., Watanabe, R., Hashimoto, T. & Nakajima, K. The Arabidopsis RWP-RK protein RKD4 triggers gene expression and pattern formation in early embryogenesis. Curr Biol 21, 1277–1281, doi:10.1016/j.cub.2011.07.001 (2011).

11 Ong-Abdullah, M. et al. Loss of Karma transposon methylation underlies the mantled somaclonal variant of oil palm. Nature 525, 533–537, doi:10.1038/nature15365 (2015).

12 Wibowo, A. et al. Partial maintenance of organ-specific epigenetic marks during plant asexual reproduction leads to heritable phenotypic variation. Proc Natl Acad Sci U S A 115, E9145–E9152, doi:10.1073/pnas.1805371115 (2018).

13 Jiang, C. et al. Regenerant Arabidopsis lineages display a distinct genome-wide spectrum of mutations conferring variant phenotypes. Curr Biol 21, 1385–1390, doi:10.1016/j.cub.2011.07.002 (2011).

14 Miyao, A. et al. Molecular spectrum of somaclonal variation in regenerated rice revealed by whole-genome sequencing. Plant Cell Physiol 53, 256–264, doi:10.1093/pcp/pcr172 (2012).

15 Han, Z. et al. Heritable Epigenomic Changes to the Maize Methylome Resulting from Tissue Culture. Genetics 209, 983–995, doi:10.1534/genetics.118.300987 (2018).

16 Stroud, H. et al. Plants regenerated from tissue culture contain stable epigenome changes in rice. Elife 2, e00354, doi:10.7554/eLife.00354 (2013).

17 Rivas-San Vicente, M. & Plasencia, J. Salicylic acid beyond defence: its role in plant growth and development. J Exp Bot 62, 3321–3338, doi:10.1093/jxb/err031 (2011).

18 Aerts, N., de Bruijn, S., van Mourik, H., Angenent, G. C. & van Dijk, A. D. J. Comparative analysis of binding patterns of MADS-domain proteins in Arabidopsis thaliana. BMC Plant Biol 18, 131, doi:10.1186/s12870-018-1348-8 (2018).

19 Mara, C. D., Huang, T. & Irish, V. F. The Arabidopsis floral homeotic proteins APETALA3 and PISTILLATA negatively regulate the BANQUO genes implicated in light signaling. Plant Cell 22, 690–702, doi:10.1105/tpc.109.065946 (2010).

20 Barwale, U. B. & Widholm, J. M. Somaclonal variation in plants regenerated from cultures of soybean. Plant Cell Rep 6, 365–368, doi:10.1007/BF00269562 (1987).

21 Etienne, H. & Bertrand, B. Somaclonal variation in Coffea arabica: effects of genotype and embryogenic cell suspension age on frequency and phenotype of variants. Tree Physiol 23, 419–426, doi:10.1093/treephys/23.6.419 (2003).

22 Tedeschi, F., Rizzo, P., Rutten, T., Altschmied, L. & Baumlein, H. RWP-RK domain-containing transcription factors control cell differentiation during female gametophyte development in Arabidopsis. New Phytol 213, 1909–1924, doi:10.1111/nph.14293 (2017).

23 Chwialkowska, K., Nowakowska, U., Mroziewicz, A., Szarejko, I. & Kwasniewski, M. Water-deficiency conditions differently modulate the methylome of roots and leaves in barley (Hordeum vulgare L.). J Exp Bot 67, 1109–1121, doi:10.1093/jxb/erv552 (2016).

24 Ferreira, L. J., Azevedo, V., Maroco, J., Oliveira, M. M. & Santos, A. P. Salt Tolerant and Sensitive Rice Varieties Display Differential Methylome Flexibility under Salt Stress. PLoS One 10, e0124060, doi:10.1371/journal.pone.0124060 (2015).

25 Seymour, D. K., Koenig, D., Hagmann, J., Becker, C. & Weigel, D. Evolution of DNA methylation patterns in the Brassicaceae is driven by differences in genome organization. PLoS Genet 10, e1004785, doi:10.1371/journal.pgen.1004785 (2014).

26 Vining, K. J. et al. Dynamic DNA cytosine methylation in the Populus trichocarpa genome: tissue-level variation and relationship to gene expression. BMC Genomics 13, 27, doi:10.1186/1471-2164-13-27 (2012).

27 Widman, N., Feng, S., Jacobsen, S. E. & Pellegrini, M. Epigenetic differences between shoots and roots in Arabidopsis reveals tissue-specific regulation. Epigenetics 9, 236–242, doi:10.4161/epi.26869 (2014).

28 Huang, Y. et al. Polycomb-dependent differential chromatin compartmentalization determines gene coregulation in Arabidopsis. Genome Res, doi:10.1101/gr.273771.120 (2021).

29 Lupianez, D. G., Spielmann, M. & Mundlos, S. Breaking TADs: How Alterations of Chromatin Domains Result in Disease. Trends Genet 32, 225–237, doi:10.1016/j.tig.2016.01.003 (2016).

30 Zheng, H. & Xie, W. The role of 3D genome organization in development and cell differentiation. Nat Rev Mol Cell Biol 20, 535–550, doi:10.1038/s41580-019-0132-4 (2019).

31 Kawakatsu, T. et al. Epigenomic Diversity in a Global Collection of Arabidopsis thaliana Accessions. Cell 166, 492–505, doi:10.1016/j.cell.2016.06.044 (2016).

32 Dubin, M. J. et al. DNA methylation in Arabidopsis has a genetic basis and shows evidence of local adaptation. Elife 4, e05255, doi:10.7554/eLife.05255 (2015).

33 Peng, Y., Yang, J., Li, X. & Zhang, Y. Salicylic Acid: Biosynthesis and Signaling. Annu Rev Plant Biol 72, 761–791, doi:10.1146/annurev-arplant-081320-092855 (2021).

34 Thomas, H. & Howarth, C. J. Five ways to stay green. J Exp Bot 51 Spec No, 329–337, doi:10.1093/jexbot/51.suppl_1.329 (2000).

35 Yoo, S. C. et al. Quantitative trait loci associated with functional stay-green SNU-SG1 in rice. Mol Cells 24, 83–94 (2007).

36 Spano, G. et al. Physiological characterization of ‘stay green’ mutants in durum wheat. J Exp Bot 54, 1415–1420, doi:10.1093/jxb/erg150 (2003).

37 Choi, J., Strickler, S. R. & Richards, E. J. Loss of CRWN Nuclear Proteins Induces Cell Death and Salicylic Acid Defense Signaling. Plant Physiol 179, 1315–1329, doi:10.1104/pp.18.01020 (2019).

38 Hu, B. et al. Plant lamin-like proteins mediate chromatin tethering at the nuclear periphery. Genome Biol 20, 87, doi:10.1186/s13059-019-1694-3 (2019).

39 Reinders, J. et al. Compromised stability of DNA methylation and transposon immobilization in mosaic Arabidopsis epigenomes. Genes & Development 23, 939–950, doi:10.1101/gad.524609 (2009).

40 Johannes, F. et al. Assessing the Impact of Transgenerational Epigenetic Variation on Complex Traits. PLoS Genetics 5, e1000530–e1000530, doi:10.1371/journal.pgen.1000530 (2009).

41 Ji, L. et al. TET-mediated epimutagenesis of the Arabidopsis thaliana methylome. Nature Communications 9, 895, doi:10.1038/s41467-018-03289-7 (2018).

42 Gallego-Bartolome, J. et al. Targeted DNA demethylation of the Arabidopsis genome using the human TET1 catalytic domain. Proc Natl Acad Sci U S A 115, E2125–E2134, doi:10.1073/pnas.1716945115 (2018).

43 Cortijo, S., Wardenaar, R., Colome-Tatche, M., Johannes, F. & Colot, V. Genome-wide analysis of DNA methylation in Arabidopsis using MeDIP-chip. Methods Mol Biol 1112, 125–149, doi:10.1007/978-1-62703-773-0_9 (2014).

44 Antunez-Sanchez, J. et al. A new role for histone demethylases in the maintenance of plant genome integrity. Elife 9, doi:10.7554/eLife.58533 (2020).

45 Borges, F., Calarco, J. P. & Martienssen, R. A. Reprogramming the epigenome in Arabidopsis pollen. Cold Spring Harb Symp Quant Biol 77, 1–5, doi:10.1101/sqb.2013.77.014969 (2012).

46 Calarco, J. P. et al. Reprogramming of DNA methylation in pollen guides epigenetic inheritance via small RNA. Cell 151, 194–205, doi:10.1016/j.cell.2012.09.001 (2012).

47 Papareddy, R. K. et al. Chromatin regulates expression of small RNAs to help maintain transposon methylome homeostasis in Arabidopsis. Genome Biol 21, 251, doi:10.1186/s13059-020-02163-4 (2020).

48 Braybrook, S. A. et al. Genes directly regulated by LEAFY COTYLEDON2 provide insight into the control of embryo maturation and somatic embryogenesis. Proc Natl Acad Sci U S A 103, 3468–3473, doi:10.1073/pnas.0511331103 (2006).

49 Curtis, M. D. & Grossniklaus, U. A gateway cloning vector set for high-throughput functional analysis of genes in planta. Plant Physiol 133, 462–469, doi:10.1104/pp.103.027979 (2003).

50 Shimada, T. L., Shimada, T. & Hara-Nishimura, I. A rapid and non-destructive screenable marker, FAST, for identifying transformed seeds of Arabidopsis thaliana. Plant J 61, 519–528, doi:10.1111/j.1365-313X.2009.04060.x (2010).

51 Krizek, B. A. & Meyerowitz, E. M. The Arabidopsis homeotic genes APETALA3 and PISTILLATA are sufficient to provide the B class organ identity function. Development 122, 11–22, doi:10.1242/dev.122.1.11 (1996).

52 Defraia, C. T., Schmelz, E. A. & Mou, Z. A rapid biosensor-based method for quantification of free and glucose-conjugated salicylic acid. Plant Methods 4, 28, doi:10.1186/1746-4811-4-28 (2008).

53 Abreu, M. E. & Munne-Bosch, S. Salicylic acid deficiency in NahG transgenic lines and sid2 mutants increases seed yield in the annual plant Arabidopsis thaliana. J Exp Bot 60, 1261–1271, doi:10.1093/jxb/ern363 (2009).

54 Koren, S. et al. Canu: scalable and accurate long-read assembly via adaptive k-mer weighting and repeat separation. Genome Res 27, 722–736, doi:10.1101/gr.215087.116 (2017).

55 Linthorst, J., Hulsman, M., Holstege, H. & Reinders, M. Scalable multi whole-genome alignment using recursive exact matching. bioRxiv, doi:10.1101/022715 (2015).

56 Fiddes, I. T. et al. Comparative Annotation Toolkit (CAT)-simultaneous clade and personal genome annotation. Genome Res 28, 1029–1038, doi:10.1101/gr.233460.117 (2018).

57 Du, Z., Zhou, X., Ling, Y., Zhang, Z. & Su, Z. agriGO: a GO analysis toolkit for the agricultural community. Nucleic Acids Res 38, W64–70, doi:10.1093/nar/gkq310 (2010).

58 Chen, S., Zhou, Y., Chen, Y. & Gu, J. fastp: an ultra-fast all-in-one FASTQ preprocessor. Bioinformatics 34, i884–i890, doi:10.1093/bioinformatics/bty560 (2018).

59 Langmead, B. & Salzberg, S. L. Fast gapped-read alignment with Bowtie 2. Nat Methods 9, 357–359, doi:10.1038/nmeth.1923 (2012).

60 Li, H. et al. The Sequence Alignment/Map format and SAMtools. Bioinformatics 25, 2078–2079, doi:10.1093/bioinformatics/btp352 (2009).

61 Sun, H. & Schneeberger, K. SHOREmap v3.0: fast and accurate identification of causal mutations from forward genetic screens. Methods Mol Biol 1284, 381–395, doi:10.1007/978-1-4939-2444-8_19 (2015).

62 Andrews, S. & Krueger, F. FastQC: a quality control tool for high throughput sequence data., 2010).

63 Bolger, A. M., Lohse, M. & Usadel, B. Trimmomatic: a flexible trimmer for Illumina sequence data. Bioinformatics 30, 2114–2120, doi:10.1093/bioinformatics/btu170 (2014).

64 Pedersen, B. S., Eyring, K., S., D., Yang, I. V. & Schwartz, D. A. Fast and accurate alignment of long bisulfite-seq reads. (2014).

65 toolkit, P. Picard toolkit. Broad Institute, GitHub repository (2019).

66 Barturen, G., Rueda, A., Oliver, J. L. & Hackenberg, M. MethylExtract: High-Quality methylation maps and SNV calling from whole genome bisulfite sequencing data. F1000Res 2, 217, doi:10.12688/f1000research.2-217.v2 (2013).

67 Hüther, P. et al. MethylScore, a pipeline for accurate and context-aware identification of differentially methylated regions from population-scale plant WGBS data. bioRxiv, 2022.2001.2006.475031, doi:10.1101/2022.01.06.475031 (2022).

68 Thorvaldsdottir, H., Robinson, J. T. & Mesirov, J. P. Integrative Genomics Viewer (IGV): high-performance genomics data visualization and exploration. Brief Bioinform 14, 178–192, doi:10.1093/bib/bbs017 (2013).

69 Wickham, H. ggplot2: Elegant Graphics for Data Analysis. (Springer-Verlag New York, 2016).

70 eulerr: Area-proportional Euler and Venn diagrams with ellipses (2019).

71 pheatmap: Pretty heatmaps [Software] (2015).

